# Sorting droplets into many outlets

**DOI:** 10.1101/2021.06.03.446983

**Authors:** Saurabh Vyawahare, Michael Brundage, Aleksandra Kijac, Michael Gutierrez, Martina de Geus, Supriyo Sinha, Andrew Homyk

## Abstract

Droplet microfluidics is a commercially successful technology, widely used in single cell sequencing and droplet PCR. Combining droplet making with droplet sorting has also been demonstrated, but so far found limited use, partly due to difficulties in scaling manufacture with injection molded plastics. We introduce a droplet sorting system with several new elements, including: 1) an electrode design combining metallic and ionic liquid parts, 2) a modular, multi-sorting fluidic design with features for keeping inter-droplet distances constant, 3) using timing parameters calculated from fluorescence or scatter signal triggers to precisely actuate dozens of sorting electrodes, 4) droplet collection techniques, including ability to collect a single droplet, and 5) a new emulsion breaking method to collect aqueous samples for downstream analysis. We use these technologies to build a fluorescence based cell sorter that sorts with high purity. We also show that these microfluidic designs can be translated into injection molded thermoplastic, suitable for industrial production. Finally, we tally the advantages and limitations of these devices.

## 1 Introduction

Emulsion droplets can be isolated, highly parallel, picoliter-scale reactor vessels. ^1^ Uniformly sized droplets, made with microfluidics, are extensively used in single cell analysis and droplet PCR, with several companies making and selling such systems. Microfluidic droplet making was first introduced in the early 2000s, ^2,3^ droplet sorting in the mid 2000s, ^4^ droplet digital PCR in early 2010s, ^5^ and methods for droplet single cell sequencing in mid 2010s. ^6,7^

Microfluidic droplet sorting is possible by many methods, but perhaps most easily accomplished by dielectrophoresis. ^8^ Aqueous drops have a higher dielectric constant compared to the surrounding carrier fluid, and an electric field gradient, applied at the right time, can pull a drop into a side channel. Thus, a drop identified via fluorescence or other means to contain a feature of interest – a cell or molecule – can be pulled out for further analysis. ^9–11^

A process uniquely suited for inexpensive, large volume manufacturing of microfluidic chips and cartridges is injection molding. While droplet making had an easy transition from elastomeric PDMS (polydimethylsiloxane) laboratory demonstrations to commercialized injection molded plastic devices, droplet sorting still presents severe manufacturability challenges, owing to the need for thick electrodes, also referred to as 3D electrodes. We describe in this paper a microfluidic design, along with associated systems and methods, to sort droplets into tens of sorting outlets with a new electrode geometry. Previously, devices made using PDMS elastomer with a few sorting junctions have been described. ^12^ We show that our design goes beyond that to tens of junctions, sorts at higher droplet rates, and crucially, is translatable into injection molded COC (cyclic-olefin copolymer) plastic.

## 2 Design

Our system can be configured as a cell sorting device, consisting of a microfluidic chip, optical apparatus including lasers, filters, detectors; mechanical apparatus consisting of chip manifolds and holders for collection; electronic apparatus including sorting circuits and signal processing circuits.

An example of a microfluidic chip design having 4 sort junctions with parts labelled, and a built chip example having 17 sort junctions is shown in Fig. 1. We will describe the parts in the sections that follow. Details of the fabrication process are available in S2 (Device Fabrication)†.

**Fig. 1.**
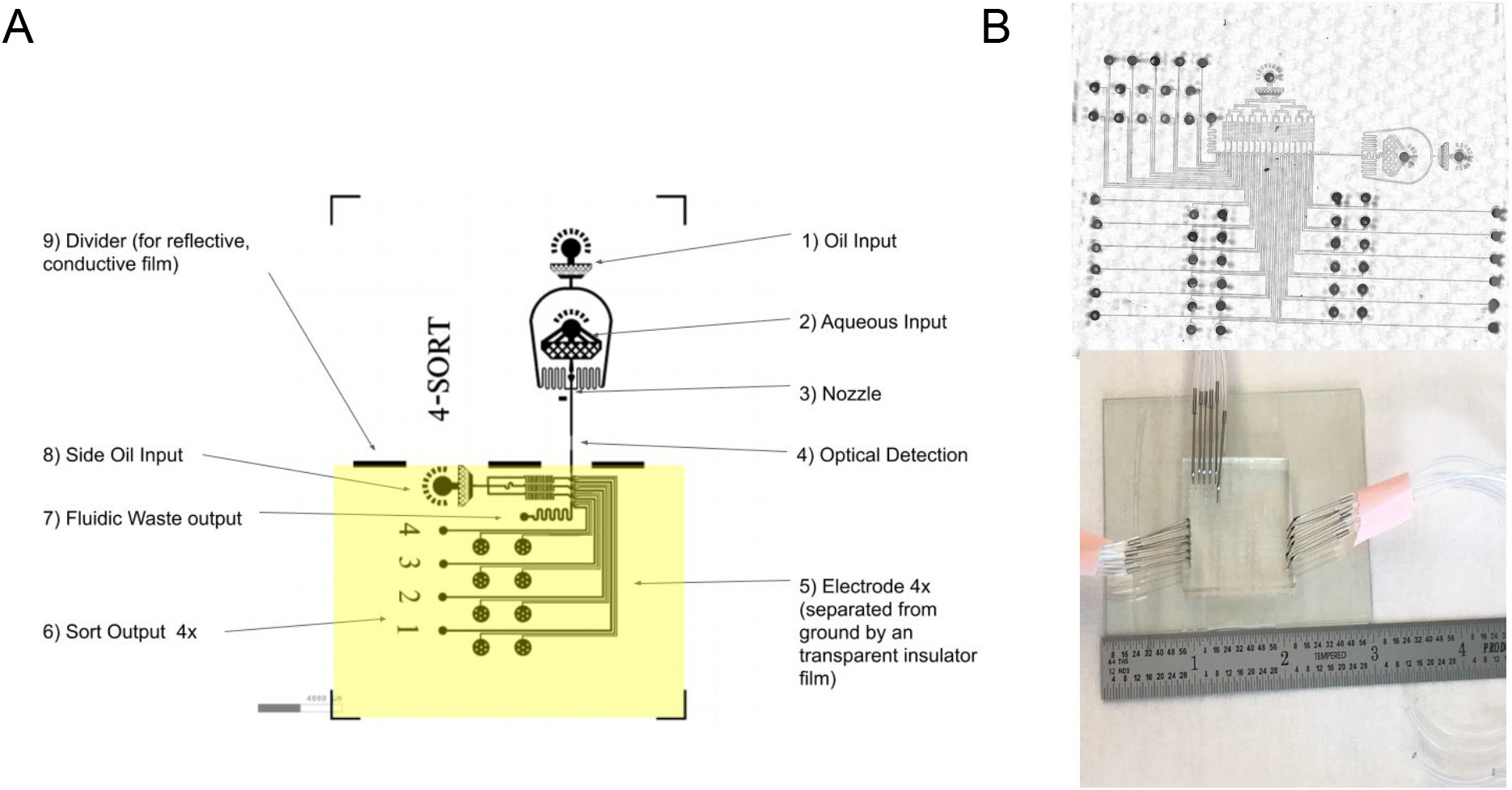
Microfluidic chip design (A) 4 sorting junction design with inputs, outputs, and other parts labelled (B) Stitched mosaic image of a 17-sort device made in PDMS. This device has a high density of features. The port holes have been punched out. Shown below is a zoomed out picture of the same device with some of the tubing attached and a ruler for scale.

### 2.1 Droplet Making

The microfluidic design starts with a flow focusing geometry, where an inert fluorinated oil and an aqueous stream meet at a nozzle to produce droplets (Fig. 2). ^13^ We use only two inlets - one each for oil and aqueous media. For fluorinated oil we used HFE-7500 (S1 Materials†). Aqueous media was distilled water or a buffer solution. The flow rates are selected to keep flow in the dripping regime, producing uniform size droplets (typically 2-4 *μ*l/min for aqueous, and 40-60 *μ*l/min for oil, using syringe pumps). Droplet production rates in our system ranges from a few hundred to 4 kHz. At higher rates we see droplet shearing and non-uniform droplet size. Droplet diameter varies from 40 to 70 micron, depending on geometry of nozzle and flow rates. ^14,15^

**Fig. 2.**
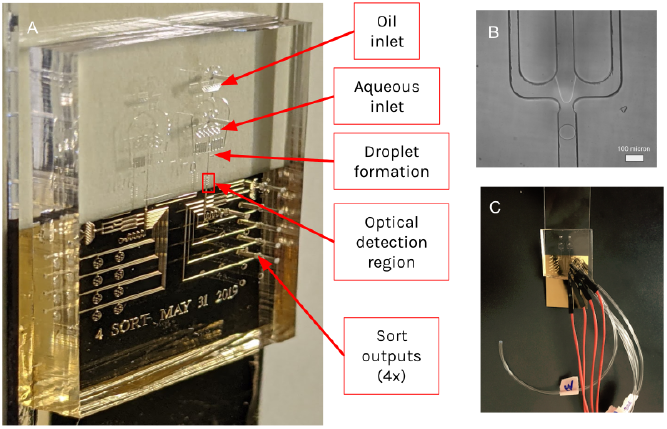
Examples of sorting devices (A) A 4-sort chip with some regions labelled. Drops formed at the nozzle flow down into the optical detection region. Using detected signals, a desired droplet may be sorted downstream into any one of the four sorting junctions the device has. Two devices are on the chip, with the unlabelled device a mirror image of the labelled one. (B) Drop production in a parallel flow geometry (in this case a plastic COC plastic chip) at a nozzle opening into a 100 micron wide channel. (C) A 4-sort chip with connecting tubes and wires. The chip is bonded to a low autofluorescence 75 mm × 25 mm glass slide coated with gold, as well as an insulating thin film of PDMS.

### 2.2 Optical Detection

The drops flow into a narrowing region where they can be probed by lasers. The overall optical schematic is shown in Fig. 3. The optical detection system consists of a set of lasers, cameras and fluorescence/scattered light detectors that provide information on what cell type is flowing through, along with information on the droplet position and velocity. Our current configuration consists of four lasers (405 nm, 488 nm, 532 nm, 647 nm), two cameras imaging from above and below the chip, and many optical detectors (typically photomultiplier tubes or photodiodes). Each of these is placed behind a chain of optical filters (Fig. 3A).

**Fig. 3.**
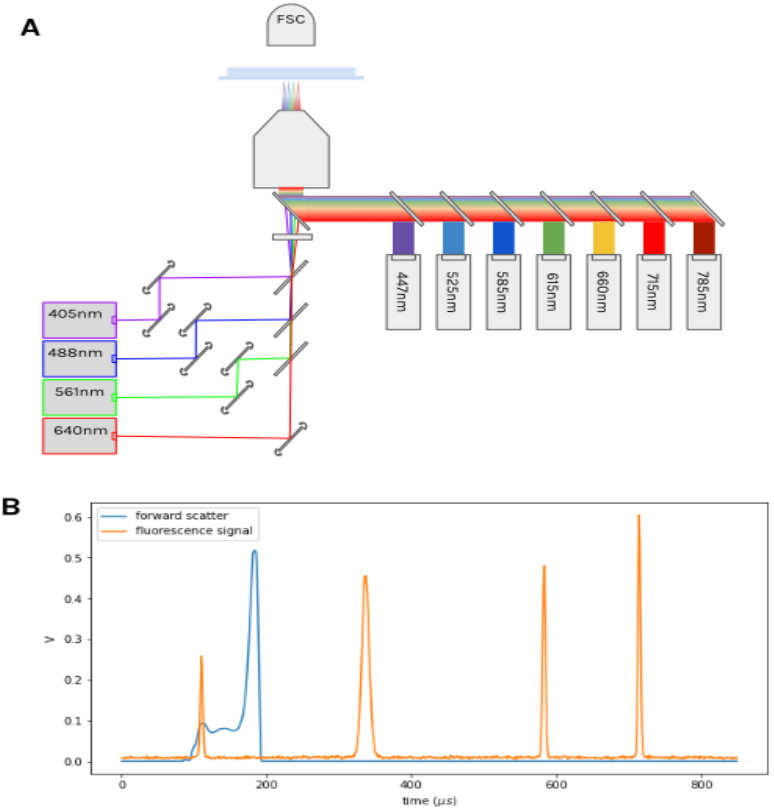
Optics (A) The optical apparatus consists of four excitation lasers, seven emission detectors, and a forward scatter detector as depicted. The emission channels are time-multiplexed to resolve each of the four excitation lasers on the single detector, producing a total of 21 excitation-emission channels. The excitation and emission path are combined using a mirror with a small masked aperture to route the low-numerical aperture excitation beam lines to the sample, while letting the high-numerical aperture fluorescence path pass to the series of PMT detectors (B) An example signal from the forward scatter detector and a single emission detector showing the four time-separated signals from each excitation laser. The forward scatter signal is dominated by scatter from the droplet itself and is useful in measuring flow rate, droplet size and frequency.

The channel height is 30 micron, and channel width is 100 micron, narrowing out to 50 micron in the optical probing region (Fig. 4). The height and width were chosen as a compromise between the fluidics and optics requirements; for optics we prefer confinement, but for fluidics an overly confined channel causes droplet shearing and breakup. ^16^

**Fig. 4.**
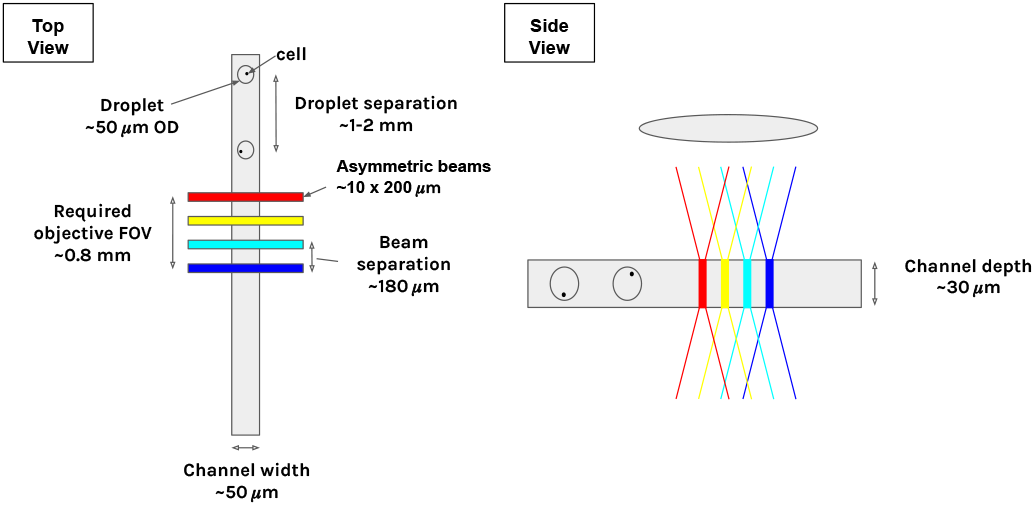
Optical probing region. 4 lasers, each shaped into an asymmetric 10 × 200 micron beam with a Powell lens, are focused in a row at the fluidic channel midplane. The size of the beam along both axes was chosen to produce near-uniform illumination across the channel cross-section. Drops with cells move through the 50 micron × 30 micron channel, and across the laser lines generating fluorescence and scatter signals.

The laser beams are shaped into a rectangular cross sectional shape for uniform illumination, and arranged in a row perpendicular to the channel, separated by ~ 180 micron. The fluorescence signals can be demultiplexed by dividing the time interval into segments that represent the time a drop spends in the vicinity of a laser. The signals can be further corrected to compensate for spillover of dye fluorescence into multiple channels, a procedure known as compensation in flow cytometry. Compared to fluorescence, scatter signals from cells are challenging to measure in our system because the droplet scatter signal overwhelms them (Fig. 3B). However, drop scatter signals are excellent for detecting drops and calculating timing from laser spot to a sort junction. More details on optoelectronics is available in ESI S3 (Optoelectronics and Software)†.

Cells or beads fill a drop randomly, with the filling distribution approximating Poisson statistics. We use concentrations such that the average filling ratio is less than 1 in 10, reducing doublets that could confound measurements. Since a cell or bead can be located anywhere in the drop, and have a velocity relative to the drop, there is a certain level of dispersion in the signals. The signal to noise quality can be improved using more confined channels, as we do in this paper, or correcting using a two color normalization approach. ^10^

### 2.3 Sorting Region

After the detection region, the drops flow into a sorting region which consists of many sorting junctions. Each sorting junction contains a sorting electrode, a sort output, and an additional oil input we refer to as side oil (Fig. 5). The side oil supplies extra oil to compensate for the loss of oil to the sorting channel (each designed to divert 30% of the main flow). Without this addition, drops would get closer to each other in the main channel, eventually colliding with each other, and making it impossible to predict timing parameters needed to sort into the downstream sort junctions.

**Fig. 5.**
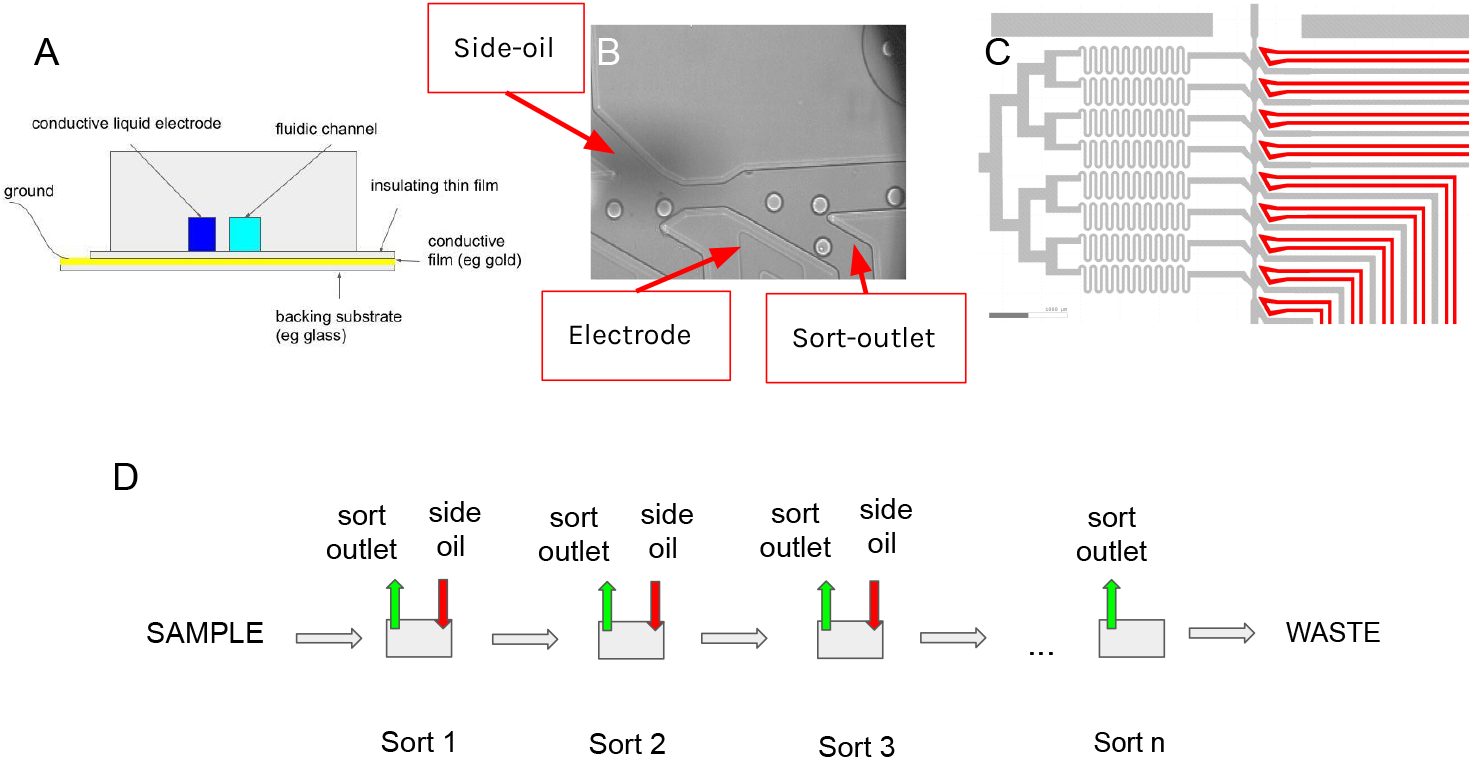
Sorting region (A) (not to scale) Diagram of the cross sectional view of a PDMS chip, showing the different layers. (B) A cell being sorted into a sort channel. The ionic liquid electrode is faintly visible. The main channel is 100 micron wide, and constricts to 50 micron just before a junction, in order to better separate drops. (C) Design of serpentine side oil channels with high fluidic resistance. These side oil channels resupply the oil lost into each sorting junction from the main channel. Also shown are the electrodes. (D) Diagram of the modular sort units, each with a sort channel and an extra oil channel (side oil). n of these units can be combined to make a n-sort device. The last sort junction does not need an extra oil supply as droplet distances do not need to be constant after all sorting is done.

We also include a constriction before each sort junction. This centers the drop, causes the drop to move faster, separating it from the following drop, and making it easier to sort.

To design the sorting region, we assumed that the fluidic network can be modelled like an electric circuit ^17^ downstream of the nozzle. In other words, we ignored any two phase related complications, and assumed laminar flow conditions. An additional simplification was made by using high fluidic resistance channels for side oil. They can then be modelled as an ideal “current” source. These side-oil channels compensate for the loss of oil, while minimizing any perturbation to flow conditions in the main channel. The design is shown in Fig. 5C where the serpentine channels are used to increase the fluidic resistance. Each side oil channel supplies the 30% oil lost in the preceding sort channel, thus keeping the volumetric flow rate in the main channel constant, which in turn keeps inter-droplet distances constant.

With the assumptions mentioned above, and given the flow rates and fixed channel heights (height determined by the fabrication process), a linear set of algebraic equations can be written down for the fluidic network, and solved for the resistances of the sort outputs. Finally, these resistances are converted into appropriate width and length for each sort output.

We found that there is considerable leeway in the design parameters, that even without precise resistances or accounting for two phase flows, devices can work for a wide range of flow rates. It is also possible to use fluidic simulation to design each side oil channel with precise resistances tuned to a particular operating pressure. However, we instead choose the simpler path of making all side oil channel resistances constant and large, relative to other resistances.

### 2.4 Electrode

Previously, sorting has been done by creating an electrode and ground channel close to the fluidic channel, filling both with low-melt solder, silver paste or salt water. ^18,19^ But the use of low melt solder/silver paste is inconvenient because of the difficulties in handling the material, while salt water can evaporate leaving behind salt crystals. Instead, our design uses an ionic liquid (1-ethyl-3-methylimidazolium tetrafluoroborate), a type of liquid salt. Ionic liquids are a large class of liquids, and one can choose a particular liquid to suit material chemistry and process conditions. ^20^

Ionic liquids do not evaporate, and these electrodes can be used for a long time (months in our experience). Liquid flow makes them easy to fill and unfill, and gives them self-healing properties. It should be noted that the electrode design will work with low melt solder, silver paste, galium or salt water without much change; however, we find the use of ionic liquid wires to be more convenient.

We made early designs using the two-electrode configurations (ground and active), both filled with ionic liquid. Later designs were further improved by removing the ground channel and using a metallic (gold) film as a common ground plane for all the ionic liquid electrodes. The electrodes were designed to be 30-50 micron away from the sort channel and filled with ionic liquid. The metallic ground plane is separated from the liquid electrode by a transparent, insulating film (PDMS or plastic), typically 20-50 um in thickness.

We did electromagnetic simulations of the electrode force fields (in Comsol): due to the ground film, the forces are directed in the plane of the chip - sideways towards the electrode (Fig. 6). This allows us to use lower voltages to sort, compared to the two electrode configuration.

**Fig. 6.**
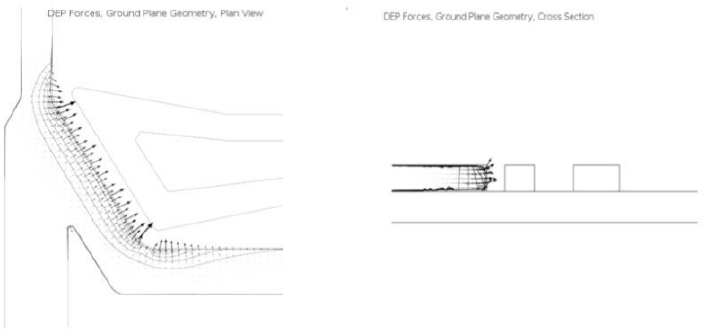
Simulation of the force field on a point particle shown in top and side cross sectional view. The electrode geometry consists of an ionic liquid filled active electrode and a ground film electrode, separated by a thin insulating film. The force is directed sideways and in the plane of the channel, a consequence of having a ground plane.

It is also possible to use ITO or silver nanowires in place of gold, if transmissivity, instead of reflectivity, is needed.

This new type of ionic liquid-metallic electrode enormously simplifies the connections required, and makes for compact designs, especially if many sort junctions are needed. The ground film also neutralizes stray charges, and serves as a mirror, allowing observation of drops when illuminated using an infra-red LED (infra-red does not interfere with fluorescence measurements). In our designs, the gold film covers only the electrode region and not the optical detection region, as optical transparency is needed for laser probing. The simplest way to do this was to use a half-coated glass slide, with gold covering only the sorting region (Fig. 2)

## 3 Device Operation

### 3.1 Sort Timing

The timing for sorting involves detecting a signal from the drop as it passes through the detection region and predicting the time it takes to arrive at a sorting junction. This detection signal may be in the form of fluorescence or scatter from the drop or its contents. In our devices, the time interval between a drop traversing from the laser spot to a particular sorting junction ranges from a few milliseconds to hundreds of milliseconds, depending on the distance and flow rates.

To be able to sort it is essential for the flow of droplets to be uniform with low dispersion in arrival time to a sorting junction. Using videos, we measured the time of arrival of drops at a sorting junction from a fixed starting line in a 9-sort device as shown in Fig. 7. The sharp peaks imply that droplet arrival time is predictable, constant over time, and the same for every drop due to the laminar nature of flow in microchannels. Once the table of timing information is known, a single trigger measurement made upstream gives us sufficient information to be able to sort that drop into *any* sort outlet reliably, provided the flow rates are constant and there are no clogs or other changes to channel dimensions.

**Fig. 7.**
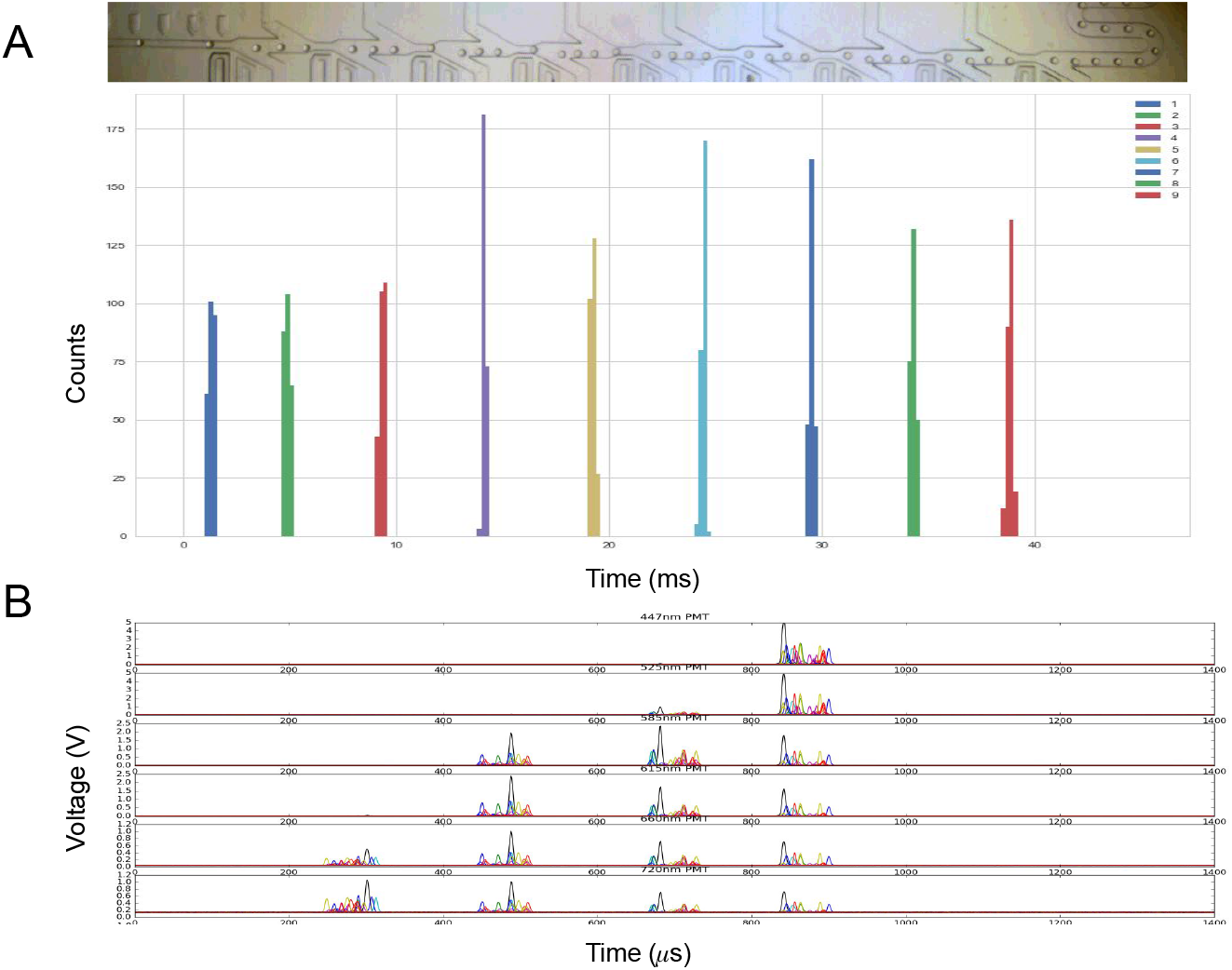
Timing drops (A) Video of drops flowing through a 9-sort device was used to compute the time of arrival of 800 drops at sort junctions. The timing is predictable without much dispersion. Histogram bin width is 500 *μ*s. (B) Fluorescence readout from rainbow beads in drops as they move across the laser line in 6 channels. The beads can be located anywhere in the drop and this results in dispersion; however, fluorescence due to each laser can be separated out in time. We are using scatter from the drop as it crosses the first laser to trigger the start of an event and collect fluorescence. We also collect a few pre-trigger signals.

To pull a chosen drop into a specific sort outlet, we apply a high voltage AC pulse (±500-800V, 10-15 kHz, 1 ms long) to that sorting electrode, exactly when a drop just enters the sorting junction.

For identification of drop contents, we also have to do a temporal/spatial separation of fluorescence signals as the drops move across laser lines. This way, fluorescence measured in a channel can be attributed to a particular laser. As shown in Fig. 7B, there is some jitter in the signal, as the bead or cell may be located anywhere in the drop. But it is possible to separate out the signals using a time window and attribute it to the correct laser-fluorophore pair.

We found that sorting a drop causes a slight speed up of the drop following it. This is because as the drop enters the sort channel, it blocks oil from flowing through, ^21^ and this increases the volumetric flow rate in the main channel, speeding up the following drop slightly. In practice, for fast flow rates, many sorting junctions, and sorting of only a small fraction of all drops, this effect did not impede our ability to sort accurately, despite not making any corrections for it.

We also find that the faster the flow, the more predictable the droplet timing, presumably because faster flow implies less time for diffusive dispersion of droplet velocity. However, for any given geometry, there is a speed limit beyond which a drop will shear into smaller droplets.

### 3.2 Scaling Number of Sort Junctions

Sort units can be stacked consecutively to produce a desired number of sort outlets (Fig. 5). While the sort junction is geometrically the same near the main flow channel, each sort outlet to such a junction needs to have its resistance designed so that the flows are the same. Fig. 8 shows the modular nature of the system, with sort outlets ranging from 4 to 17 (also see Video S1, S2, S3, S4†). We believe that this is the highest number of sorting junctions in an individual emulsion sorting design demonstrated in literature so far.

**Fig. 8.**
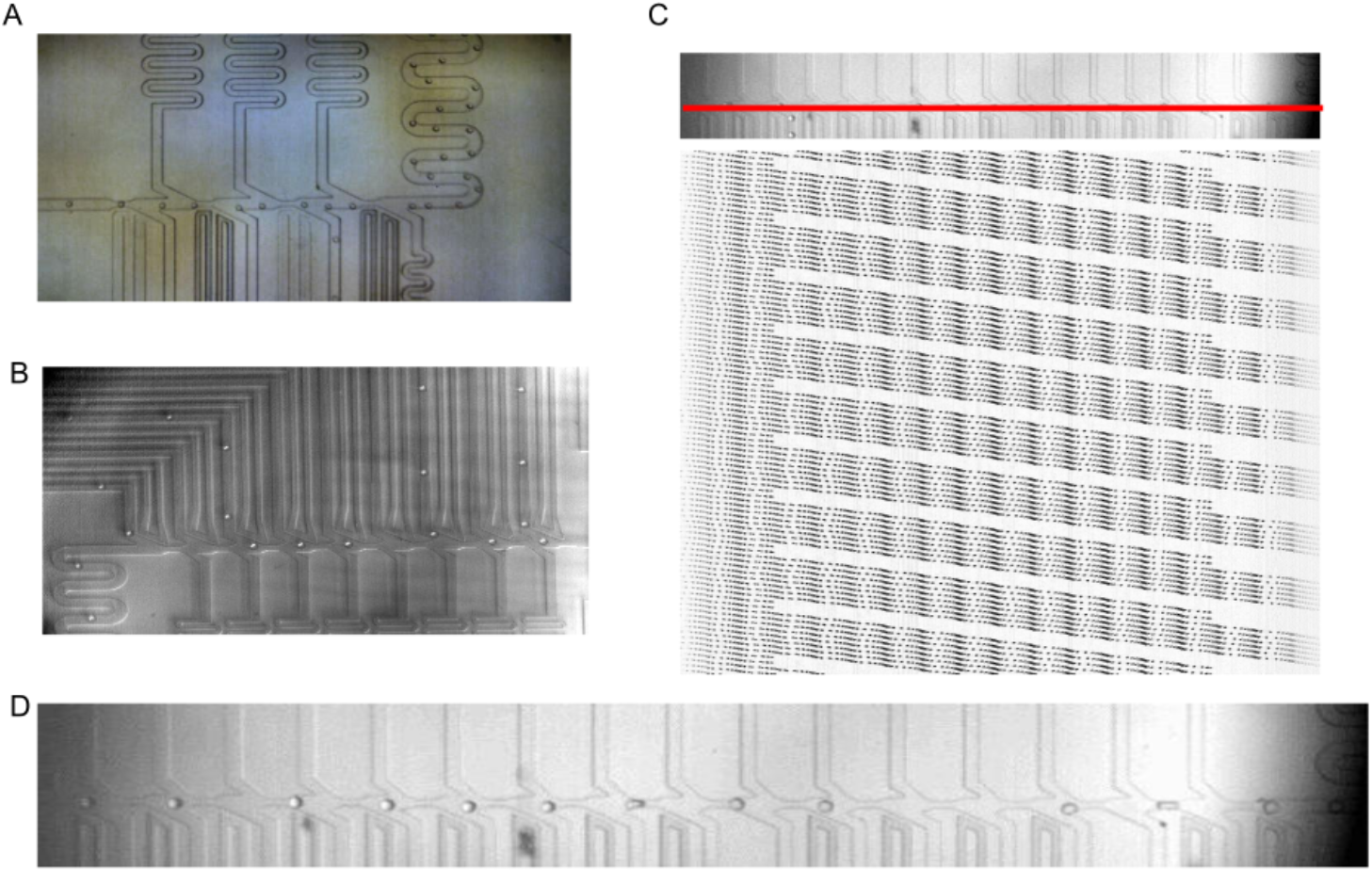
Modular Sorting Junctions: Sorting junctions can be combined to produce a device with desired number of sort outlets. The main channel is 100 micron in width in all cases and constricts to 50 micron before a sorting unit. (A) 4-sort design with two ionic liquid electrodes (ground and active) for each sort junction (B) 9-sort design, each junction having a single single active electrode. A gold film serves as common ground for all electrodes (C) Kymograph showing continuous sorting of two drops each into junction 3 and 15 in a 17-sort device. Drops are flowing at 225/second. The horizontal dark bands are the sorting junctions. The slanted lines are individual droplets. Where the line abruptly stops, a drop has been sorted. Faster droplet velocity causes the lines to become non-continuous and dotted. The red line across the main channel was used to generate the kymograph (D) A snapshot of the sorting region from a 17-sort device (also see Video S2†).

Using a line pixel intensity scan, and combining all the lines over a stack of video frames into a single image, we can visualize drops moving in time and space. Due to the refractive index change at the drop-oil boundary, it is easy to identify a drop. If the line scan is chosen along the main flow channel, the drops show up as continuous lines or streaks, terminating if they get sorted. Faster flow or lower camera frame rate results in fuzzier, dotted lines. Fig. 8C demonstrates such a trace, called a kymograph, on a 17-sort junction device. Two drops are being sorted in junction 3, the next two in junction 15, followed by six drops not sorted. This sorting pattern is continuously running (Video S2†). These kymographs were made using FIJI software. ^22^

The predictability in timing lets us expand the number of sort junctions, limited only by the large flow rate needed (each sort junction diverts part of the flow, needing resupply for compensation), camera field of view, and tubing connections.

Using videos, we can calculate the droplet velocities and trace the path of drops (Video S1†). This is shown in Fig. 9, where the paths of drops are overlaid on top of each other, with a color map of the velocities. If the sort outlets have appropriately designed fluid resistances, the flow pattern at each sort junction should look like the others.

**Fig. 9.**
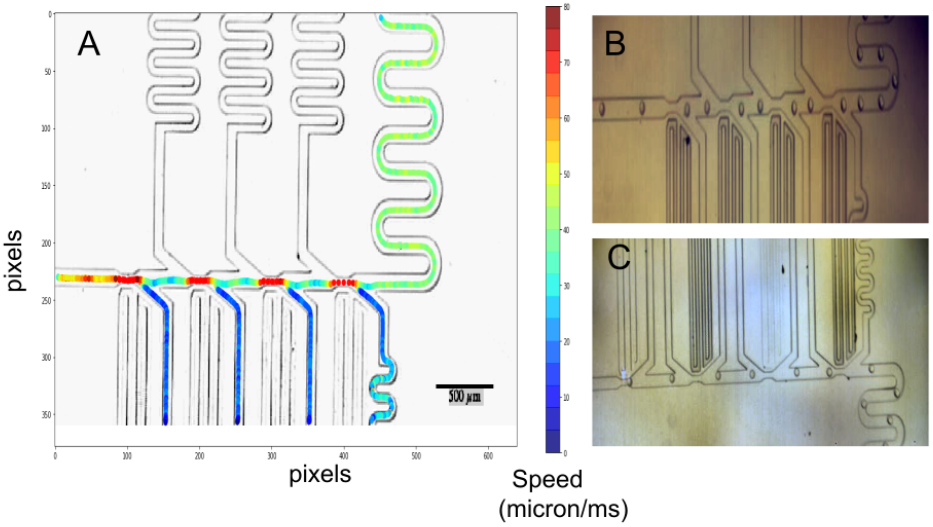
Drops flowing through sorting junctions (A) Using videos, the path and velocity of 200 drops were overlaid on a 4-sort design. Drops were being sorted in 1234W1234W1234W... pattern where the number is the sort junction and W is waste. Notice the similarity of flow at each sorting junction (B),(C) The side oil supply may be arranged to be on the same side or the opposite side to the sort channels. The opposite side configuration is more compact compared to the same side design, but either may be used.

As demonstrated in Fig. 9B and C, the side oil supply may be arranged to be on the same or opposite side to the sort channels. The opposite side configuration is more compact, and the one we used for most of our designs.

### 3.3 Droplet Flow in Tubes

If predictable recovery of intact drops was required, we found it essential to use narrow bore tubing at the device sorting outlets. This is because in larger bore tubes, drops can get trapped at the side walls in slower moving fluid and won’t be swept out. This leads to droplets colliding, and with every collision comes with a probability of coalescence; eventually plugs form, moving at unpredictable speeds (Fig. 10A). But with narrow bore tubing, we were able to predictably recover even a single sorted drop, as demonstrated in Fig. 10. In narrow bore tubing, there is space only for one drop in the lateral direction. Drops are exposed to the fastest flowing oil at the center of the tube, and swept away with that oil flow.

**Fig. 10.**
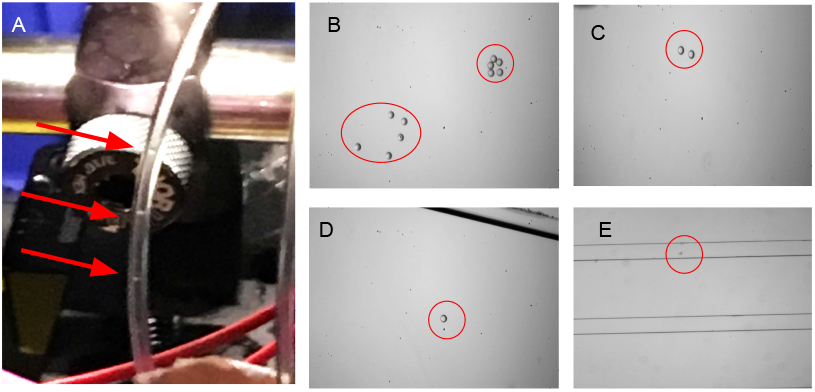
Effect of tubing on sorted drops (A) Sorted drops coalesce to form plugs moving at unpredictable rates in a large bore tubing (500 micron here). The plugs can be observed as the tube is transparent. (B) (C) (D) Narrow bore tubing prevents coalescence. Here 125 micron inner bore was used. 5, 2 or 1 drops were sorted for every 1000 drops, with a droplet rate of 350/s. If we collect fluid from the sort outlet, we see a pulse of droplets every 3-4 seconds. The drops are <100 pl, and the oil collected is less than 5 *μ*l. (E) We connected the two sort outlets of a 2-sort to another chip with two long channels, using narrow bore tubing. The channels were imaged on a microscope. Single drops were sorted into one of the sort junctions, and then observed to flow by several seconds later in the channel.

The volume of a 50 micron drop is only about 65 picoliter, so precise collection of a small number of drops opens up the possibility of performing picoliter scale combinatorial biochemistry where drops may be dropped into wells and combined with other drops, each containing cells, beads or chemicals. ^23^ The presence of inert oil prevents these drops from evaporating rapidly. In air, they would only last a few seconds before completely evaporating.

### 3.4 Emulsion Breaking

For many downstream processes, the droplet emulsion needs to be broken, and the aqueous phase collected for subsequent steps. We developed a method involving corona discharge to break the emulsion. Alternative ways to break the emulsion involve the use of an anti-static discharge gun. ^24^ Standard chemical methods involving perfluoro-octanol (PFO) may also be used, but the chemical can interfere with downstream processes, and be difficult to completely remove from the solution.

In the sort collection tube, collected droplets float to the top of the oil. Emulsions collected, for example in a centrifuge tube, were broken by exposing the tube to a corona discharge machine (BD-20AC Laboratory Corona Treater, Electro Technic Inc, Chicago, IL) for a few seconds. Once the emulsion was broken, the aqueous phase could be pipetted out and used in further downstream processes.

The effects of using corona discharge for breaking of the emulsion on cells was measured using the GM12878 cell line. To separate the effects of putting the cells through a microfluidic device and/or sorting from the effects of corona discharge on cells, AF647 labeled GM12878 cells and unlabeled GM12878 cells were used to make an emulsion by shaking the cell-oil mixture. The emulsion was left on ice for 30-45 min. Subsequently corona discharge was used to break the emulsion and the aqueous phase was extracted along with the cells. We measured cell viability using Trypan blue on the Countess (Thermofisher, Waltham, MA) as well as using Nucleocounter (Chemometec, Denmark), following the manufacturer’s instructions. There was n’t a significant difference in viability between the control GM12878 cell sample and the cells that were recovered from the emulsions using the corona discharge treatment (Table 1). The cell samples were also analyzed using flow cytometry on the Attune Acoustic Focusing Flow Cytometer (Life Technologies, Thermo Fisher Scientific), and no changes in scatter or fluorescence were observed between the corona discharge treated and untreated cells (data not shown).

**Table 1.**
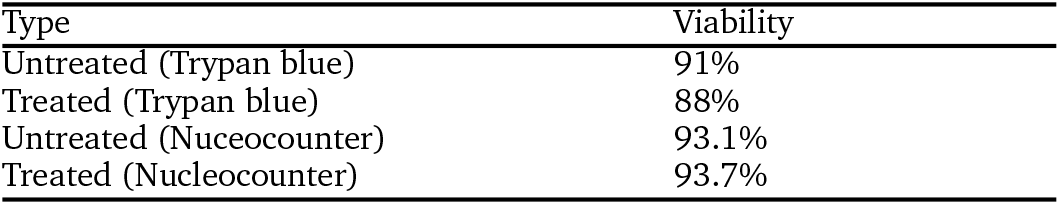
GM12878 cell viability on treatment with corona discharge

| Type | Viability |
| --- | --- |
| Untreated (Trypan blue) | 91% |
| Treated (Trypan blue) | 88% |
| Untreated (Nucleocounter) | 93.1% |
| Treated (Nucleocounter) | 93.7% |

## 4 Sorting Beads and Cells

To test system performance, we used 2-sort, 4-sort, and 9-sort devices to sort calibration beads and labelled cells (Video S4†). Sorted drops flow down to the sorted outlet port, and then via microbore tubing (125 micron ID PEEK tubing) into a sort collection tube. Fig. 12 shows examples of fluorescent beads sorted, collected, and observed under a microscope

**Fig. 11.**
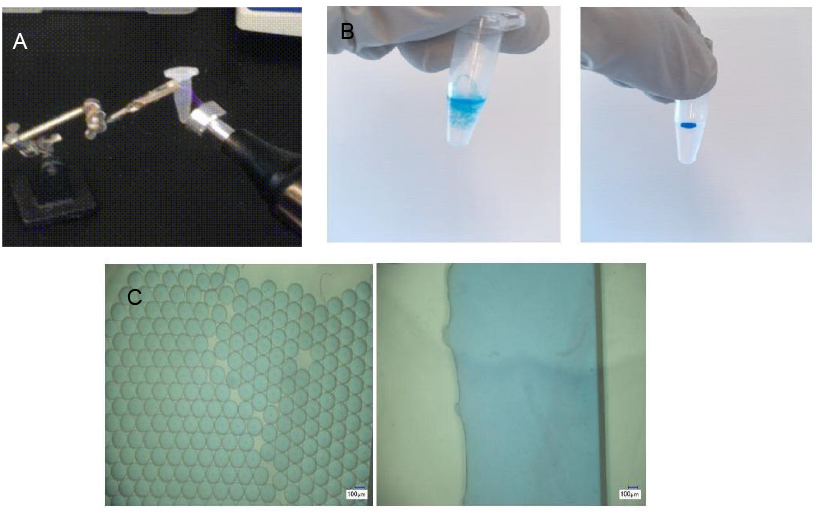
Emulsion breaking using a corona discharge machine. (A) Corona discharge wand brought close to a centrifuge tube for emulsion breaking (B) Before and after image of an emulsion on application of corona discharge. (C) Before and after photos of the emulsion breaking in a plastic haemocytometer.

**Fig. 12.**
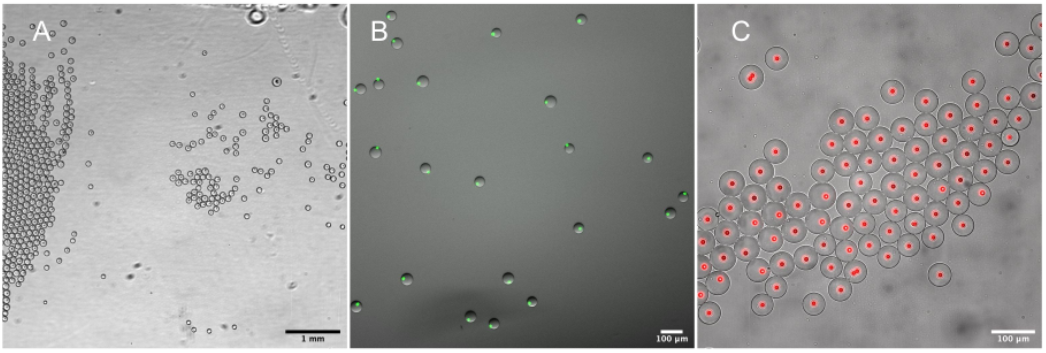
Sorting fluorescent beads (A) Drops from a sort experiment collected in a haemocytometer. (B) Fluorescence (Alexa 447), pseudo color, overlaid on a brightfield image. In this case, the sorting was done on a 4-sort plastic chip. (C) Sorted drops with Alexa 647 fluorescent beads (pseudo color), in this case collected from a 4-sort PDMS device.

Use of narrow bore tubing from the syringe to the aqueous inlet was found to be essential to sweep cells or beads into the chip. In this regard, we also found it useful to use small magnetic stir bars to gently agitate the syringe contents, reducing sedimentation. Cells were also cooled in the syringe using a home-built syringe cooling sleeve contraption.

For small droplet numbers and quick testing, a filling chamber like a haemocytometer may be used to collect the flow from an outlet tubing to observe under a microscope (Fig. 12A).

Purity measurements on sorted drops were made by either manual counting in a haemocytometer under a microscope or breaking the emulsion, pipetting the aqueous fraction, and running the aqueous portion on a flow cytometer.

Fig. 13A shows an experiment where red and green labelled cells were sorted in a 2-sort device. We used GM12878 cells and CD45 antibodies labelled with AF488 or PE-Cy5. The labelled cells were mixed in approximately 1:1 ratio, with each type filling about 2% of the drops. Using fluorescence signals from the two channels, we drew gates, and sorted each type into a separate sort output. These sorted cell-laden droplets were collected, the emulsion broken using corona discharge, and aqueous phase reflown in a flow cytometer along with controls. From Fig. 13A we see that the final measured purity was about 89% for cells labelled with AF-488 and 95% for cells labelled with Pe-Cy5 in this experiment.

**Fig. 13.**
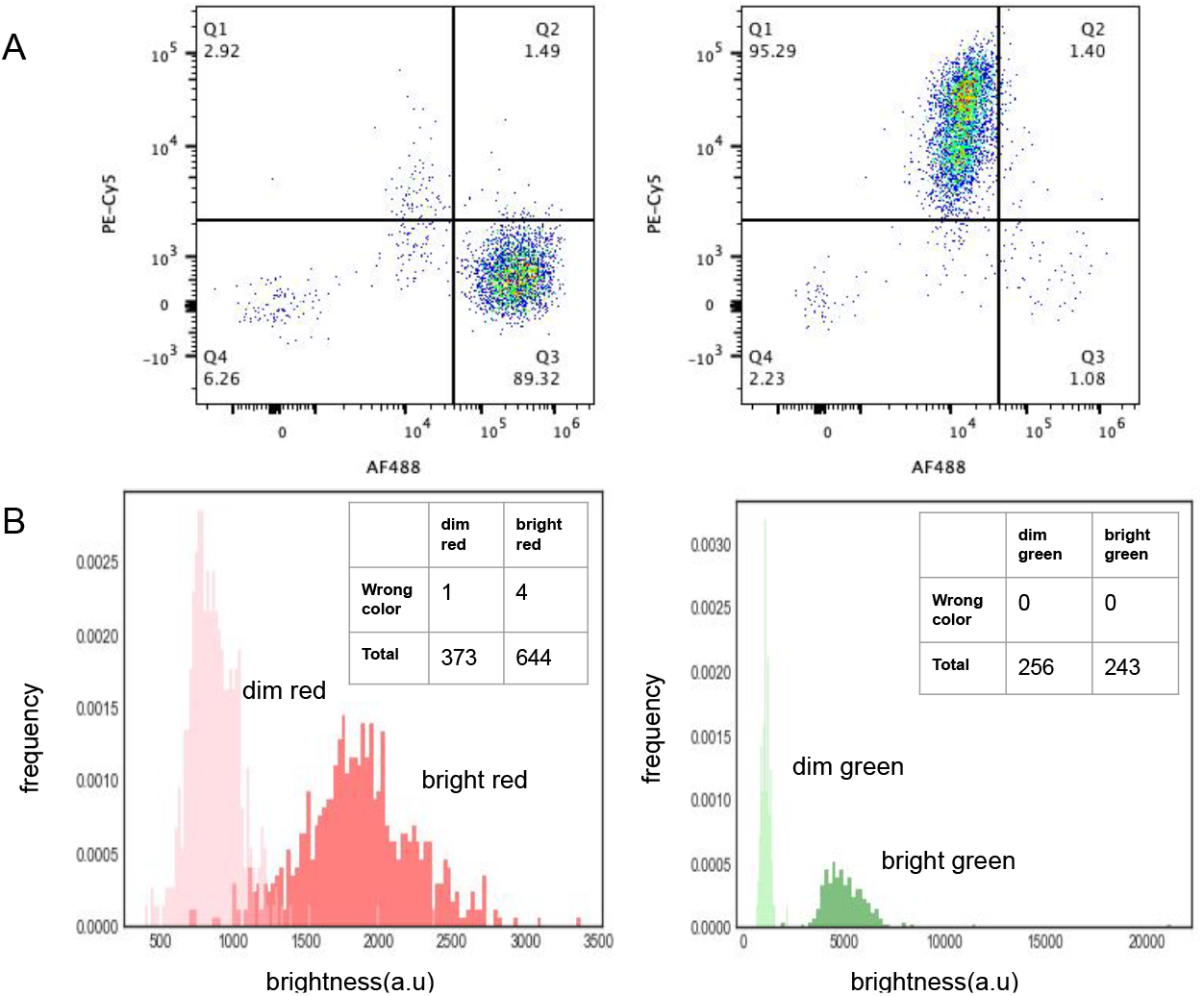
Sorting beads and cells (A) AF488 and Pe-Cy5 labelled cells were mixed in approximate 1:1 ratio and then sorted in a 2-sort device. We collected drops from each sort outlet, broke the emulsion, collected the aqueous phase, and ran it on a different flow cytometer (Attune) to quantify purity. Roughly 89% and 95% purity was obtained respectively for each sort outlet (B) Dim and bright green and red fluorescent beads were sorted on a 4-sort device, droplets collected, and their fluorescence intensity estimated from microscope images to create a frequency histogram. Each peak is from a different sort output. The ability to sort red/green is tabulated in the inset tables and shows high spectral discrimination. (Intensities measured from images are broader than the intrinsic bead fluorescence, due to the difficulty in measuring fluorescence intensities from a microscope image).

Fig. 13B shows an experiment using a 4-sort device, where drops filled with red and green beads of two different intensities (Bangslab Alexa 488 and Alexa 647 MESF 3 and 4) were sorted into each of the 4 outlets. Each bead type was present in approximately 2% drop filling ratio. The sorted drops with beads were collected from the outlet sort tubes into a haemocytometer, and the fluorescence evaluated by processing the images (Fig 12). We see the ability to separate out the dimer and brighter fluorescence in both the colors. The inset tables count the color missorts (green for red or vice versa) corresponding to each sort outlet. Overall, due to the sharp and bright fluorescence from beads, our optical detection has little difficulty in detecting the beads. Fluorescence from labelled cells tends to be broader and noisier.

Another bead sort experiment with rainbow beads (Spherotech Ultra-rainbow 5 fluorescence levels) done on a 9-sort device, with 5 of the outputs being used, showed the purity in Table 2 (as measured using a separate flow cytometer). Note that sort purity increases as the beads get dimmer; this is because the fluorescence detection system is unable to distinguish doublets, and assumes that the brighter bead is the correct one, leading to impurities in the sort outputs with higher fluorescence.

**Table 2.** Rainbow beads with 5 fluorescence levels (5 = highest, 1 = lowest) were sorted in a 9-sort device, with 5 outputs being used. Following this, the outputs were collected, emulsion broken and flown in a flow cytometer (Thermo Fisher Attune). Each level filled the drops 5-8%, resulting in a doublet rate of about 1-3%. On the flow cytometer we obtained over 500 beads in each level. Percentages may not sum to 100% due to rounding. The purity percentages show a clear trend with purity increasing as the dimmer beads are sorted. This is due to doublets being sorted into the sort channel of the highest fluorescence bead. Carryover in the flow cytometer also contributes to error.

| Type | Sort output 1 (%) | Sort output 2 (%) | Sort output 3 (%) | Sort output 4 (%) | Sort output 5 (%) |
| --- | --- | --- | --- | --- | --- |
| Level 5 | 91.4 | 0.9 | 0.2 | 0.0 | 0.0 |
| Level 4 | 2.2 | 91.3 | 0.3 | 0.0 | 0.0 |
| Level 3 | 2.4 | 2.9 | 95.0 | 1.1 | 0.1 |
| Level 2 | 2.1 | 2.7 | 2.2 | 96.0 | 1.7 |
| Level 1 | 1.9 | 2.2 | 2.3 | 2.9 | 98.1 |

In general, we were able to achieve sorting at purity that exceeded 90% in most of our experiments, with the highest purity exceeding 99.5%. The main source of error is doublets (our detection system was not designed for this task). Beyond doublets, the signal to noise of optical detection is another factor limiting purity. Other sources of errors involved incorrectly set timing parameters and cell debris clogging a channel.

We expect that the sorting precision of our droplet system to be on par or exceed that of high speed flow sorters because the fluids are moving at an order of magnitude slower velocity, and we have the ability to sort individual drops precisely.

Besides high purity sorting, it is also possible to enrich a rare population, setting parameters such that borderline cases may still be sorted. In this case, we favor yield over purity.

## 5 Plastic Devices

While PDMS is an extraordinary material for rapid prototyping and testing in a laboratory, it has shortcomings as a material for large scale industrial manufacture. ^25^ Instead, injection molded plastic is the material/method of choice for inexpensive, large volume manufacturing.

We translated the design into injection molded COC plastic with a few minor changes. ^26–28^ Fig. 14 A,B,C show examples of plastic devices. Figure 14D shows the cross-sectional schematic. Due to the resolution limitations of injection molding, features need to be more spread apart and sharp inner corners rounded. High precision machining is needed to make the masters molds. The dimensions of the channels are otherwise comparable to the PDMS devices. The fluidic lines go to the edge of the chip where port holes allow connections to tubing.

**Fig. 14.**
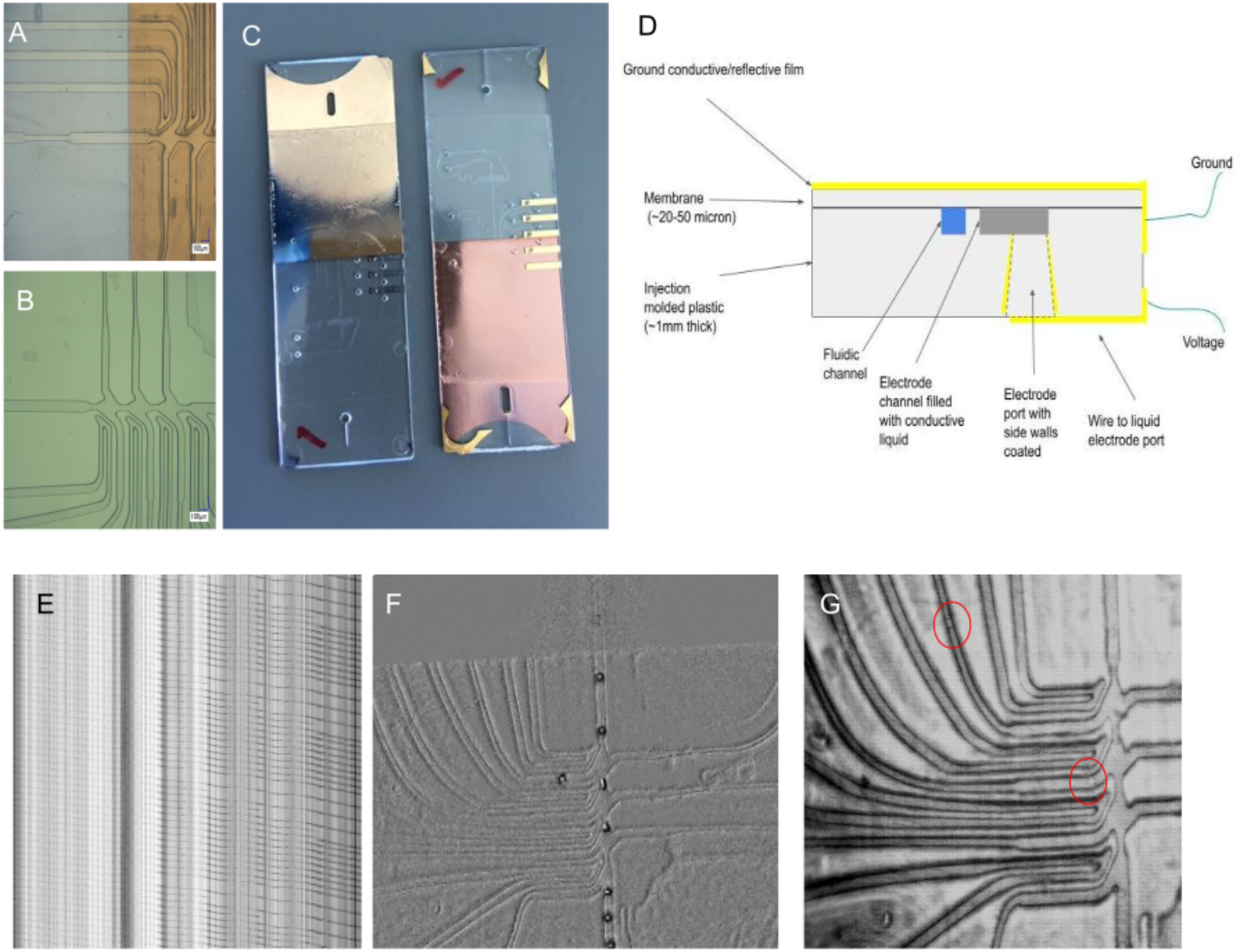
The 4-sort design was translated into injection molded COC plastic. The plastic design is almost the same as in PDMS, except that the channels are more separated from each other and have rounded corners. (A) Gold coated COC chip. The gold film serves as the ground plane for the electrodes B) similar to A) but without any gold (C) Front and back of the final device (75 × 25 mm). The gold film and the gold traces are visible. The thin membrane cover boundaries can also be observed. (D) Instead of a glass slide, we bond a thin COC membrane to make the cover as shown in the schematic image (not to scale). Gold is coated on both sides. The electrode channels are filled with ionic liquid which makes electrical contact with the gold traces (E) Kymograph of sorting in a plastic COC device in a 1×2×3×4× pattern where x = no sort, and 1, 2, 3, and 4 are sort channels. Drops were flowing at approximately 600 per second (F) A still from a video frame with the background subtracted out to show droplet sorting (Video S5†). (G) 2 drops sorted in outlets 1 and 2 in a plastic device (the roughness is due to limitations of precision machining needed to make injection molds).

We built a device with 4 sort junctions. Instead of a glass slide cover, a thin film of COC served as the channel top cover. The thinner this layer, the lower the actuation voltage. Our devices used films varying between 25 to 50 micron in thickness. Details of the fabrication process are available in ESI S2.2 (Device fabrication) †. Compared to other thermoset plastics, COC is ideal for our application due to its optical transparency, low autofluorescence and chemical inertness.

To make electrodes, we deposited a gold film on top of the membrane covering half the chip (the area with electrodes) to serve as the ground plane. On the other side, we created gold traces that lead from the ionic electrode port to the edge of the chip where we could connect to them electrically (Fig. 14D). The gold coats the inner walls of the ports allowing an electrical connection to the ionic liquid. A manifold with o-rings was used to interface with the chip and external tubing.

We were able to successfully sort using AC voltage pulses similar to ones used in PDMS devices, with peak voltage between ±500-700V (Video S5†). Fig. 14 E,F,G demonstrate examples of droplet sorting. The same kind of droplet timing used with PDMS devices is directly applicable here. Fig. 12B is a picture of droplets with beads that were sorted in a plastic device.

Plastic devices provide a scalable route to industrial manufacture. After the injection molding masters have been made, large numbers can be molded at low cost per device. With injection molding, standardization and reproducibility is more easily achieved. Hence, conversion to plastic is an important step in moving a technology from a niche demonstration to more widespread use.

## 6 Discussion

In the last decade, a whole suite of tools for manipulating droplets upstream and downstream of sorting has developed. ^29^ Microfluidic FACS (fluorescence activated cell sorting) systems ^30^ that do use emulsion droplets, ^31^ as well as those that do not use emulsion droplets ^32,33^ have been attempted. While the first FACS system used a microfluidic device, ^34^ most current FACS systems use sheath flow to position cells in a small detection cross section followed by aerosol generation at a nozzle. These aerosol drops are charged and sorted by deflection plates held at a voltage.

Compared to emulsions, aerosol generating systems can run samples at much higher droplet rates (60k/s or more), obtain sharper fluorescence signals by locating cells in a narrower probe region (typically 5-10 *μ*m), and have a mature support and service infrastructure, honed over decades this technology has been in use. These FACS systems are also larger in size and cost, have less predictability in sorting (due to detection of cells before droplet formation), are limited to 2-6 sorting outlets (due to the high speed flow and limitation imposed by plate deflectors and multichannel optical detection) and pose safety risks of biohazard aerosolization.

Both systems face difficulties in dealing with clogs and shear stresses on cells. Microfluidic droplet systems have an advantage if isolation of cells or molecules is required. Compared to droplets in air, emulsion drops evaporate much slower due to the surrounding oil, despite being only tens of picoliters in volume, making them good for viable single cell applications. Their low volume also ensures high reactant concentrations that improves yields, and their separation isolates reaction products, avoiding dilution.

In individual microfluidic devices, emulsion droplet making is limited to approximately 4 kHz at a single oil-water nozzle. At higher rates, sorting is still possible, but droplet shearing and non-uniform droplet size is observed. In our system, while we can sort at the maximum drop generation rate, the rate limiting step is in fact the signal processing limits of our current electronics/software, particularly when measuring fluorescence signals in all channels. In that case, we were limited to sorting at about 1 kHz. While not attempted in this work, it is possible to use faster electronics (at greater cost), parallelize designs, ^35,36^ or separate droplet generation from droplet sorting to obtain more throughput. ^37^

We chose a linear system with one main flow channel with sorting channels branching off in order to simplify the geometry and optical measurements, but binary systems where the main channel itself branches off into two channels, four channels, eight channels, and so on are also possible, though they would require 3D geometries to implement a similar oil resupply strategy. It is also possible to design on-chip drop collection traps, ^38^ provided only a fixed number of drops trapped is acceptable in the experiment.

Besides cell sorting, our system can be alternatively configured for high throughput screening (for antibodies, enzymes, molecules or cells) instead, where one or more sorting junctions is used in conjunction with moving well plates, to deposit individual drops into a well for further processing. This is possible due to the predictability of droplet flow through narrow bore tubing after sorting. In this case, costs can then be reduced with simpler detection systems, as finely resolved, high dimensional fluorescence data is typically not needed for most screening, and image processing in conjunction with a simpler single channel fluorescence detection is suitable. Further, COC plastic is replaceable with other thermoplastics, if high optical transparency or autofluorescence is not a concern.

## 7 Conclusions

We have designed, made, and tested a new droplet sorting microfluidic system. Two new parts introduced are an electrode combining ionic liquid and metallic films, and resupply lines to keep inter-droplet spacing constant. We were able to stack sorting junctions and make individual designs with more than a dozen sorting junctions. Sorting with this device allows precise collection of aqueous droplets, that may be only a few picoliters in volume. We also invented an emulsion breaking method, involving corona discharge, that is scalable and easy to use. Putting it all together, the system was used to sort cells and beads with high purity. These chip designs were translated to injection molded plastic devices that are inexpensive to manufacture, and offer standardization and reproducibility advantages over “artisanal” elastomeric devices made at a research lab.

Our droplet sorting technology is compatible with currently used droplet-based single cell sequencing and droplet digital PCR applications. Oil-water droplet systems, while lower in throughput than air-water systems, have an advantage if biochemical processes need to be isolated from each other. Processes like antibody generation, enzymatic screening, cell lysis, cell-cell, or cell-bead interactions. Future directions of our work may involve porting this technology to such applications, parallelizing the designs, and combining with upstream and downstream droplet processing modules that add functionality to the system.

## Supporting information

Electronic Supplementary Information

## Conflicts of interest

The authors are employees of Verily Life Sciences LLC, a subsidiary of Alphabet Inc, that has a commercial interest in this technology, and has filed for multiple patents based on this paper’s contents.

## Acknowledgements

We wish to acknowledge Holger Becker and Christian Bose of microfluidic ChipShop for plastic device making help; Trey Shoop of Polyplastics Inc. for material selection advice. From Verily: Matias Heinrich, Robbin Sneeden, Adam Walker, Dan Barrows for help with machining/injection molding; Chris Baker, Robin Simmons and John Balibalos for cytometry assistance; Alisha Knudson, Michael Hopkins for cell culture; Ninfa Sandoval, Junjia Ding, Kye Lee, Jeff Clarkson, Son Ly for microfabrication help; Xiaomi Du, Biyu Li, Licheng Xian, and Pouya Kheradpour for downstream processing and analysis; Mathew Mammen and Srinivas Hanasoge for help with plastic devices; Charlie Kim and Jess Mega for advise.

