## Supplementary material for "Sorting droplets into many outlets": Electronic Supplementary Information

### S1 Materials

We use Novec HFE-7500 (3M, Minneapolis, MN) as the inert fluorinated oil. For surfactant, we added Evagreen oil (Bio-Rad, Hercules, CA) to HFE-7500 in a 1:5 ratio. Aqueous fluid was buffer or distilled water. PBS was used with beads and RPMI media was used for tests with mammalian cells. Media was obtained from ThermFisher (Waltham, MA), microbeads were obtained from Spherotech Inc (Lake Forest, IL) and Bangslabs (Fishers, IN).

PDMS devices were made using the recommended 1:10 ratio of Sylgard, and the material was obtained from WPI (Sarasota, FL). Topaz COC pellets for injection molding and making films were obtained from PolyPlastics USA (Farmington Hills, MI). All other chemicals were obtained from Sigma Aldrich (St Louis, MO). Tubing and other fluidic components were obtained from IDEXX Inc (WestBrook, ME). Syringe pumps were obtained from KDS Scientific (Holliston, MA).

### S2 Device Fabrication

Designs were made in AutoCAD/KLayout (for PDMS) and Solidworks (for COC).

#### S2.1 PDMS Devices: Softlithography

PDMS devices were made by photolithography with SU8 3050/3035 photoresist (Kayaku Advanced Materials, Japan - formerly Microchem Inc) followed by soft lithography. Holes were punched with a coring punch (Syneo Accu Punch MP10). The devices were bonded using oxygen plasma treatment (Harrick Plasma Cleaner PDL-001-HP) to gold coated glass slides covered with a spin coated insulating layer of PDMS (2600 rpm for 1 min). Gold was coated on half the glass slide, covering only the electrode region, while leaving the optical detection region transparent.

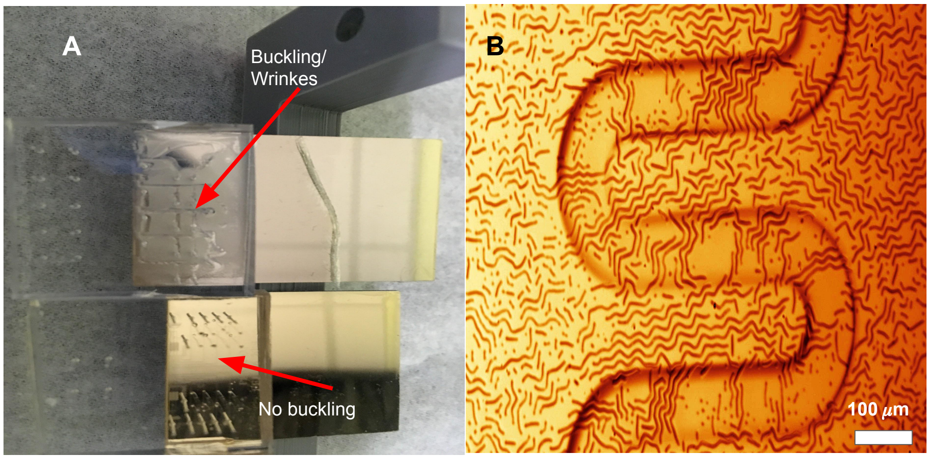


**Figure S1** PDMS wrinkles on gold (A) A comparison between PDMS devices bonded to a gold coated slide: one without a glass coating on top, and another with the coating (B) A PDMS serpentine channel 100 micron wide (part of a 2-sort device) that develops wrinkled features when a film of glass was not deposited on the gold (false color)

A small layer of silicon dioxide was deposited on top of the gold in order to improve bonding to PDMS. If we did not deposit this layer we found a wrinkling/buckling effect, and eventual delamination of the device (Fig. S1). The deposition scheme used was a layer of chromium (5 nm) to improve gold adhesion followed by gold (30 nm), and a thin layer of glass (20 nm). The device channels were not treated, but used a few days after bonding, when the hydrophilicity had worn off.

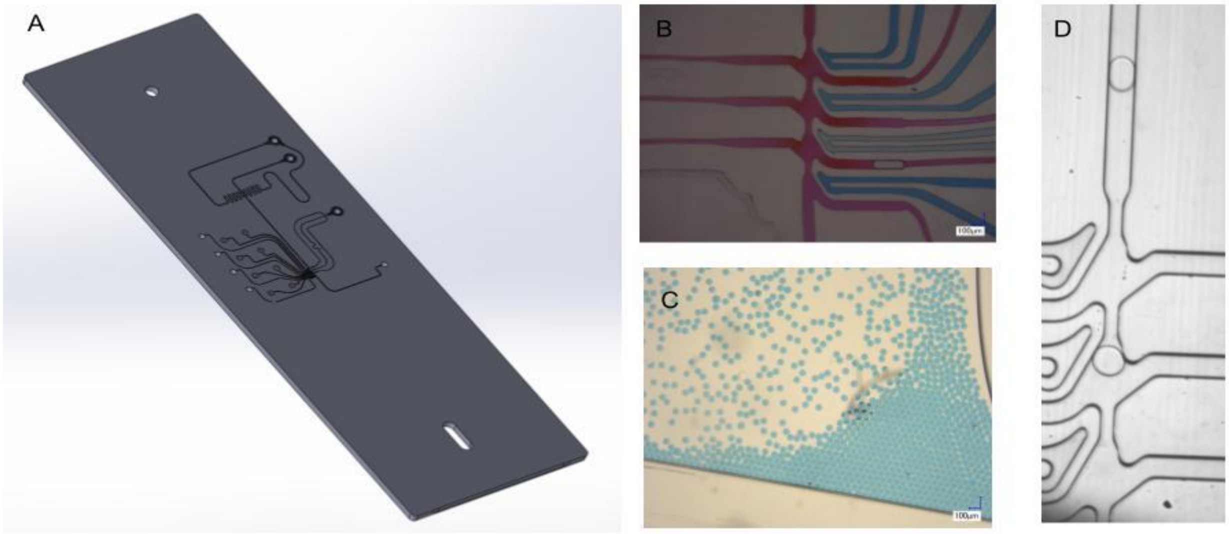


**Figure S2** Plastic Devices (A) A CAD model of the 4-sort COC plastic device. The device dimensions are 75 mm x 25 mm (B) Device filled with dyes to make flow channels and electrode channels clear. This is a view of the sorting section. The electrodes are 35 um at their closest approach to the flow channels in this picture (C) Drops produced by the device (D) Example of droplet production in a 4-sort device. The main channel is 100 micron at its widest, and the electrode is 50 micron at the nearest approach from the sort channel. A 50 micron film serves as top cover.

#### S2.2 COC Devices: Injection Molding

Plastic COC devices were made in two ways. One, externally, to our specification by microfluidics Chipshop (Jena, Germany). An off-the-shelf 50 micron COC membrane was bonded to create a cover for these devices. Two, internally, Verily Life Science's machining/injection molding lab made devices for us. The process involved precision machining an aluminum mold followed by injection molding. We used Topaz 5013L for the bulk plastic chip, and made cover membranes by dissolving COC Topaz 8007X in sec-butyl benzene at a 30% v/w concentration, and spin coating a film at 1000 rpm on a glass slide. The film was dried on a hot-plate at 100 C for a few hours. Following this, the films were released, cut, and bonded to the injection molded bulk COC chip. We used a hot press (Carver 4386) to lightly adhere the film, with a final bonding step in an oven.

We used a laser cut brass shadow mask or a photolithographic brass shadow mask (Fotofab Inc, Chicago, IL) to selectively coat gold using e-beam evaporation, both for the gold ground film on the membrane side, as well as the connections to the electrodes on the other side.

These devices are comparable in performance to PDMS devices with the exception that high power blue/UV lasers can cause the holes to develop in the thin membrane, possibly due to a small amount of energy absorption by the plastic at those wavelengths.

### S3 Opto-electronics and Software

The optical detectors include photomultiplier tubes to measure fluorescence (Hamamatsu, Japan), and SiAPD detectors (Thorlabs) to measure scattered light. Two cameras above and below the chip are used for imaging and alignment (Point Gray Grasshopper3 and Phantom Veo 640). One camera focuses on the sorting region using illumination provided by an infra-red LED, reflecting off the ground gold film and illuminating the chip. The other camera is focused on the optical detection region.

The electronics consist of three systems: analog input, digital processing, and analog output. The analog input system is a series of custom PCBAs which condition and filter the signal from the optical detectors to allow for digitization of the signal. A two stage amplification system consisting of a transimpedance amplifier and a differential amplifier convert the current output of the detectors into the appropriate voltage for the ADCs.

The gain of the PMTs is digitally adjustable via a custom PCBA with a multi-channel high resolution DAC. The fluorescence signal is digitized by analog to digital converters (ADCs) integrated on a microprocessor (Texas Instruments, Dallas, TX). The ADC's digitize the signal at 1 MSPS and 16 bits. The microprocessor's integrated voltage comparators with digitally adjustable thresholds are used to initiate digitization and timestamp events. The analog output system has individually controllable outputs for each sort junction present on the microfluidic chip. A custom PCBA takes a digital logic input, and when the logic level of the input is high, the output is a high voltage AC signal which is connected to the electrodes on the chip. The power supply for the high voltage system consists of two PS300 DC high voltage power supplies (Stanford Research Systems, Sunnyvale, CA), and a 33500B frequency generator (Agilent/ Keysight, Santa Rosa, CA) that supplies the clock signal.

#### S3.1 Control Firmware

The ADCs constantly digitize the signal from the fluorescent detectors into the microprocessor RAM using a ping-pong buffer scheme and the processor's DMA functionality to minimize processor overhead. When the forward scatter detector signal crosses the threshold of a voltage comparator, the digitization is programmed to retain a fixed number of pre-trigger samples in memory, and to fill the remainder of a fixed size sample buffer. A peak detection algorithm performs a background subtraction and determines if any fluorescent signal

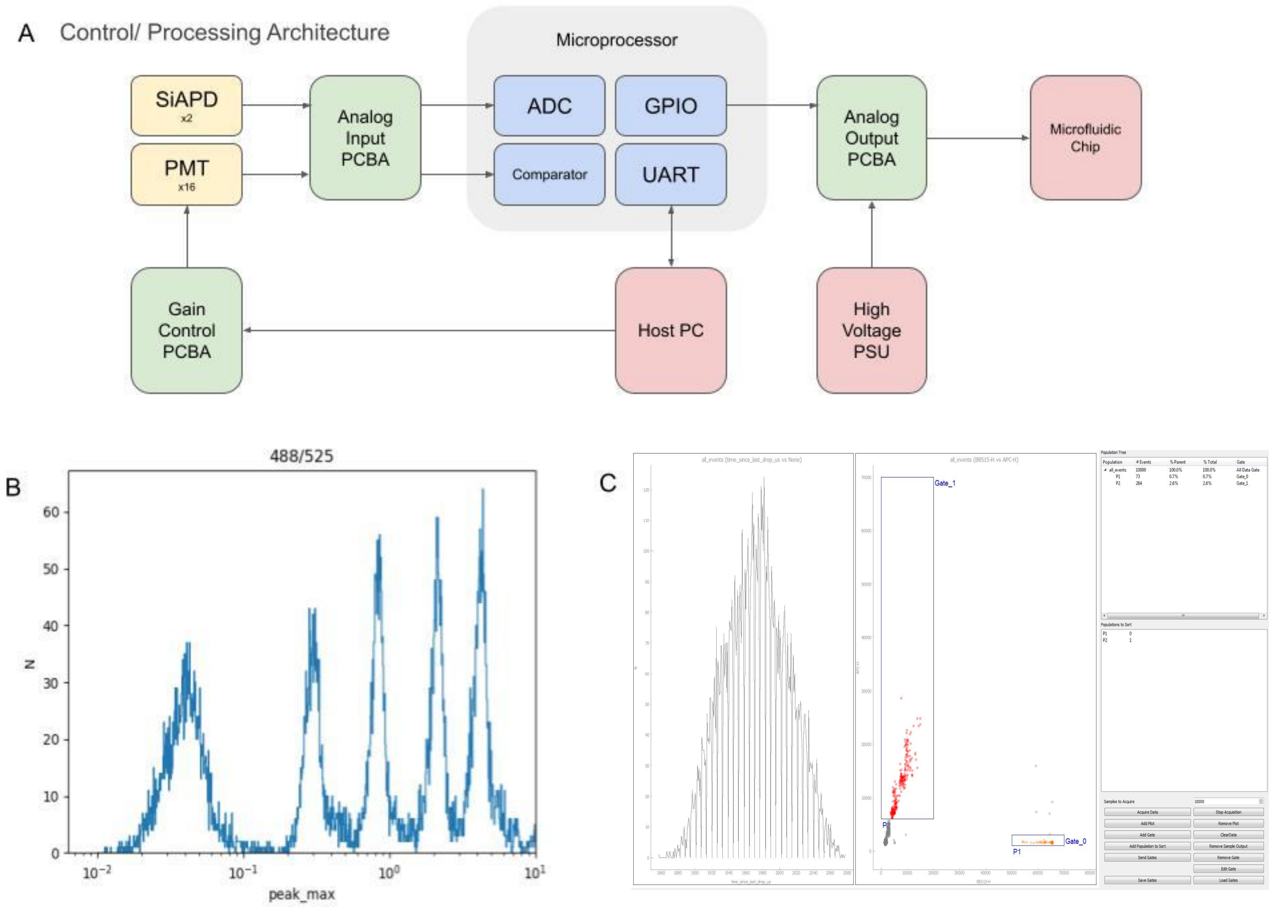


**Figure S3** Control system and software Interface (A) The overall system architecture. Analog signals from detectors are converted into digital signals. The scatter signal from a drop is used as a trigger to start acquisition. The fluorescence/scatter signal received can be used to decide whether to sort a drop into a particular output (B) We are able to measure fluorescence to a 3 log dynamic range in many channels. The figure shows fluorescence collected from a set of MESF fluorescence beads with 5 levels of fluorescence (C) The software's graphical interface enables setting the timing parameters, and selecting the gates for sorting into a particular sort channel.

was detected for each excitation/emission pair. The spatial separation of the lasers is used to demultiplex the signals originating from different excitation lasers. The peak height, peak width, and peak area are calculated for all events. The statistics from a single droplet are compared to user provided parameters (i.e. gates) and a sort/no sort determination is made. If a droplet is to be sorted, the microprocessor determines the latency until the droplet is in the correct sorting zone on the chip for the appropriate junction. At the correct time, a scheduling algorithm activates a digital output signal which is fed into the analog output system.

### S3.2 User Interface

The microprocessor sends data to and receives commands from a host PC over a serial interface. A user interface displays data and operational metrics received from the microprocessor. Users can create traditional flow cytometry plots and gates to control the sorting behavior of the system, as well as adjust thresholds and timing parameters. Datasets can be stored on the PC to be viewed later.

### S4 Videos

#### Video S1

Droplets sorting in a 11223344WW pattern, where the number is the sort output and W is waste. The approximate droplet velocity was calculated from video frames overlaid as an arrow, with some jitter. Droplet rate is approximately 90 Hz. The camera frame rate was 2500 frames per second.

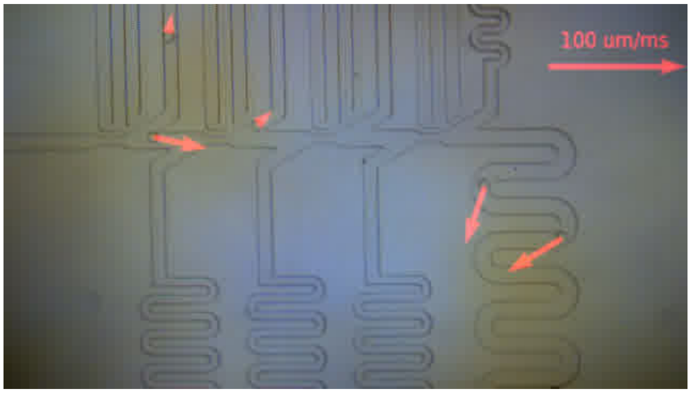


**Figure S4** Snapshot of Video S1

#### Video S2

17 junction sorting with two drops being sorted in junction 3 and 15. Drops flowing at approximately 230 per second. Original video taken at 2500 frames per second.

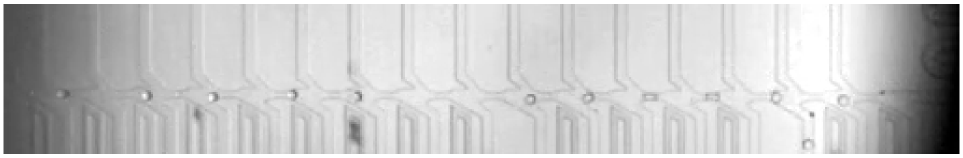

**Figure S5** Snapshot of Video S2

#### Video S3

Drops being sorted such that 1 in 3 drops go into each of the 8 sort outputs. Droplet rate is 660 Hz. Original video taken at 10000 frames per second.

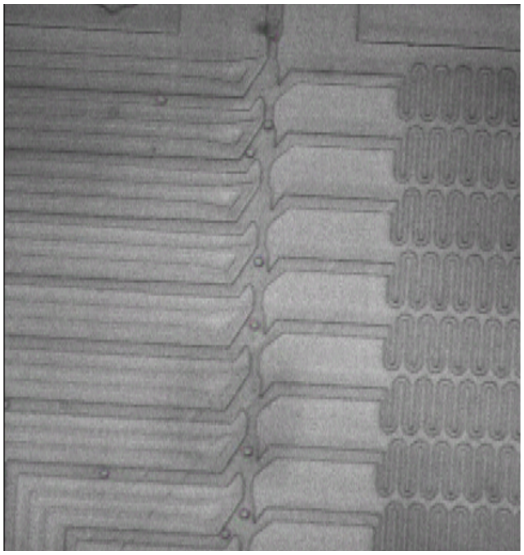

**Figure S6** Snapshot of Video S13

#### Video S4

Cells being sorted into two sort outlets based on fluorescence labelling with FITC and Pe-Cy5 dyes. Original video taken at 10000 frames per second.

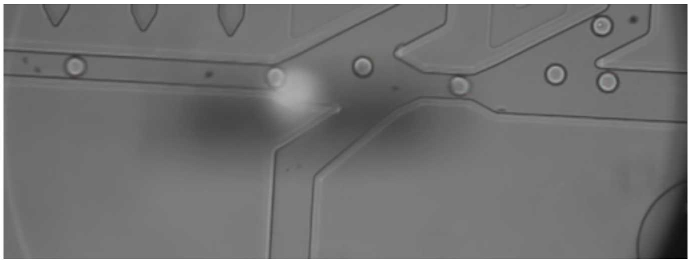

**Figure S7** Snapshot of Video S4

#### Video S5

Droplet sorting in a COC plastic 4-sort chip. The background has been subtracted to make the droplets more visible. Drops are being sorted in 1x2x3x4x order where x is a no-sort and 1,2,3,4 are the 4 sort outlets. Drop frequency was about 600 Hz. Camera frame rate is 10000 frames per second.

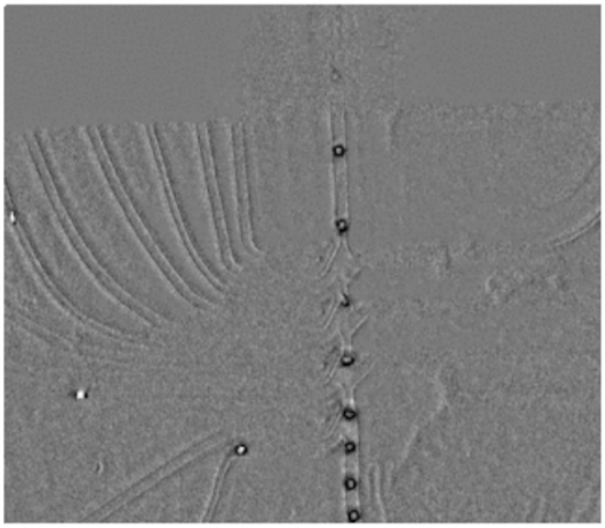

**Figure S8** Snapshot of Video S5
